# Superovulation and aging perturb oocyte-granulosa cell communication

**DOI:** 10.1101/2023.10.30.563978

**Authors:** Klaudija Daugelaite, Perrine Lacour, Ivana Winkler, Marie-Luise Koch, Anja Schneider, Nina Schneider, Alexander Tolkachov, Xuan Phuoc Nguyen, Adriana Vilkaite, Julia Rehnitz, Duncan T. Odom, Angela Goncalves

## Abstract

*In vitro* fertilization has been developed to overcome reduced fertility, which is increasingly due to a decline in reproductive cell quality during aging. Here, we quantitatively investigated the interplay between superovulation and aging in mouse oocytes and their paired granulosa cells using newly adapted isolation techniques. We tested the hypothesis that superovulation disrupts oocyte maturation, revealing the key intercellular communication pathways dysregulated by forced hormonal stimulation. We further demonstrated that granulosa cell transcriptional markers can prospectively predict an associated oocyte’s early developmental potential. By using naturally ovulated old mice as a non-stimulated reference, we showed that aging and superovulation dysregulate similar genes and interact with each other. By comparing mice and human transcriptional responses of granulosa cells, we found that age-related dysregulation of hormonal responses and cell cycle pathways was shared, though substantial divergence exists in other pathways.

**Highlights:**

- Superovulation perturbs cumulus-oocyte communication
- Granulosa cell transcription predicts superovulated oocyte quality
- Superovulation and aging non-additively perturb similar sets of genes

## Introduction

During mammalian ovulation, oocytes are released into the female reproductive tract as cumulus-oocyte-complexes (COCs) composed of one oocyte (OC) surrounded by somatic granulosa cells (GC). Mammalian reproduction relies on release of COCs containing high quality oocytes. In both humans and most other animals, fertility sharply decreases with age due to low oocyte numbers^1,2^ and poorer oocyte quality, reflecting gene expression dysregulation and/or inadequate maturation.^3–6^ In mice, oocytes from old individuals have increased aneuploidy rates,^7–9^ gene expression dysregulation,^7,10^ and lower developmental potential^11^. In response to age-related fertility drop in humans,^12–14^ assisted reproductive technologies (ART) have been developed, which use hormonal stimulation (superovulation) to collect oocytes, followed by *in vitro* fertilization (IVF) to generate embryos.^15,16^ In humans, ART procedures lack efficiency and have a strikingly low ratio of eggs to live births, which further decreases for older patients.^17–19^

A crucial element of successful IVF is the identification of high quality oocytes and embryos. Currently, most common clinical approaches select embryos based on their gross morphology, and result in a low percentage of live births.^20–22^ Less commonly used approaches include invasive preimplantation genetic testing, which is linked to adverse obstetric and neonatal outcomes^23^ leading to a growing interest in identifying alternative methods for calling oocyte quality.^24–27^ It has been speculated that an oocyte’s quality could be evaluated by transcriptional patterns in surrounding granulosa cells.^28^ Indeed, during IVF granulosa cells that could be used for non-invasive screening are routinely collected yet discarded. Identifying a reliable transcriptomic signature connecting granulosa cells to clinical outcome has remained elusive,^29^ due to variation in IVF procedures and retrospective study designs,^30^ small sample numbers,^27^ non-sensitive detection methods,^31,32^ and pooling of COCs.^33^

Current models that explain the lower quality of superovulated eggs as reflecting developmental immaturity were based on morphological and biochemical characterization,^34–37^ fertilization and implantation rates,^38–40^ and oocyte epigenetics.^41–43^ Final oocyte maturation occurs via nuclear and cytoplasmic changes,^44^ requiring correct chromosomal separation^45^ and maternal transcript remodeling,^46–48^ respectively. However, the oocyte can only mature if it establishes correct bi-directional communication with the surrounding cumulus granulosa cells.^49–54^ To better understand the developmental potential linked to the initial oocyte quality, a close investigation of ovulated COC is necessary. However, previous efforts to separate individual oocytes and granulosa cells were limited to post-mortem surgical extraction of immature cells from the ovary in mice,^55,56^ cows,^57^ monkeys,^58^ and humans.^6^

How aging shapes the response to superovulation also remains poorly understood, often due to model limitations. For instance, many studies have been limited to a single isolated cell type, either oocyte or granulosa cells.^4,59,60^ Another approach has been to compare the oocytes obtained from young and old animals following superovulation.^10,61,62^ To our knowledge, currently there are no studies that decouple the effects of superovulation and natural aging in cumulus-oocyte complexes that have been released by the ovary.

Here, we compare and quantify the effects of aging and superovulation on COCs, and demonstrate that oocyte early developmental potential can be predicted by profiling surrounding granulosa cells. To accomplish this, we employed a rigorous study design with two novel features. First, we manually isolated granulosa cells from around each oocyte for paired analysis, and, second, we obtained naturally ovulated COCs in young and old mice for comparison with superovulated COCs. Our improved study design identified the perturbations in oocyte maturation and cell-cell communication driven by hormonal stimulation of the ovary.

## Results

### Manual dissociation of cumulus-oocyte complexes into paired components

In order to simultaneously analyze oocyte and granulosa cell transcriptomes from individual COCs, we developed a novel protocol to manually separate the two while retaining their in vivo pairing information (Methods, Figure 1A, Figure S1A, Supplementary Movie S1). Using a SMART-seq2 protocol, each oocyte was sequenced as a single cell and its surrounding granulosa cells as a small bulk (Methods). For each experimental condition and cell type, we typically collected 15-31 replicates, and in 72% of these replicates both paired components successfully passed QC (Supplementary Table S1 and S2). To increase statistical power, non-paired oocytes and/or granulosa cells were included in specific analyses, as described below.

**Figure 1.**
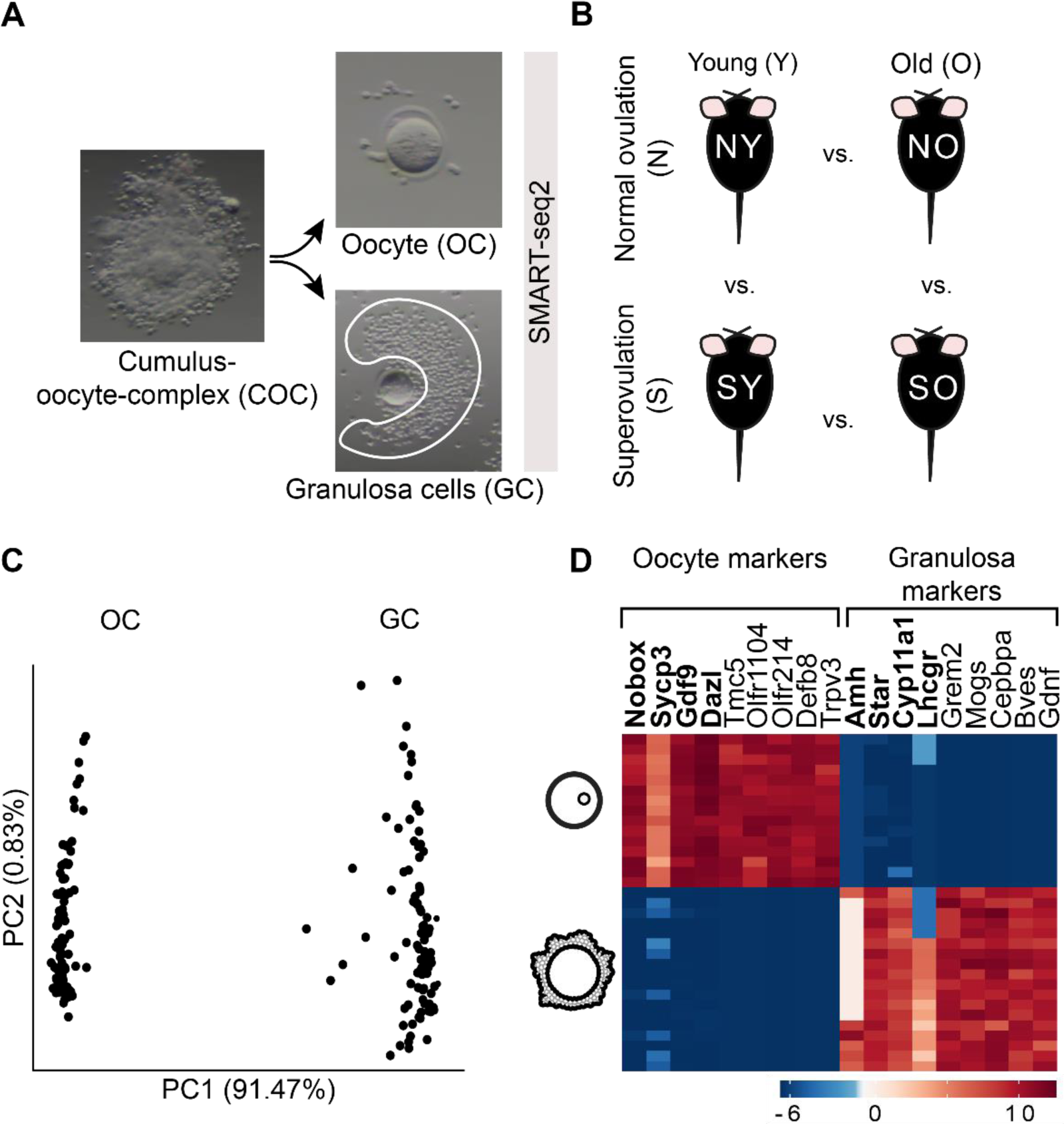
Trackably cleaving cumulus-oocyte complexes (COCs) into their two component cell types in naturally and superovulated young and old mice. (A) Manual separation of COCs into oocyte (OC) and small bulk of its granulosa cell (GC) pairs, followed by single cell sequencing using SMART-seq2 (Methods). (B) COCs were collected from naturally ovulated young (NY, n = 3), superovulated young (SY, n = 4), naturally ovulated old (NO, n = 3), and superovulated old (SO, n = 3) mice. (C) PCA representation of all OC (n = 98) and GC (n = 93) samples from all four experimental groups. (D) Expression of cell-type-specific markers in OCs and GCs. First four markers for each cell type were selected from literature, and the last five were identified from our data.

We collected paired COC transcriptomes from 3-month old (henceforth ‘young’) and 12-month old (henceforth ‘old’) mice in four conditions: naturally-ovulated young (NY), superovulated young (SY), naturally-ovulated old (NO), and superovulated old (SO) (Figure 1B). The oocytes and granulosa cells from all conditions were readily distinguished by their full transcriptomes (Figure 1C). We confirmed no cross-contamination between oocytes and granulosa cells by inspecting the transcription of previously known marker genes^58,63–67^ (Figure 1D). Exploiting the high number of replicates, we identified multiple novel oocyte- and granulosa-specific markers with similarly polarized gene expression (Figure 1D, Supplementary Table S3).

### Superovulation perturbs key COC maturation pathways

We reasoned that any developmental immaturity from superovulation should be reflected in simultaneous dysregulation of related pathways in oocytes and their associated granulosa cells. In young mice, we first compared the impact of superovulation on all available oocyte and granulosa cell transcriptomes. In oocytes, we identified 3519 genes (of 22181 genes detected) that were differentially expressed following superovulation (adjusted p-value < 0.05, 46% upregulated) (Figure 2A). As expected, many pathways critical for oocyte development were perturbed (Supplementary Table S4), including meiosis, mitochondrial metabolism, DNA damage, and hormonal response.^37,68–70^ In granulosa cells, superovulation dysregulated 2765 genes (of 25201 detected) (adjusted p-value < 0.05, 40% upregulated) (Figure 2B). Similarly to oocytes, cell-cycle, hormonal and damage response pathways were enriched in differentially expressed genes^71,72^ (Supplementary Table S4). Overall, superovulation disrupts pathways required for successful COC development.

**Figure 2.**
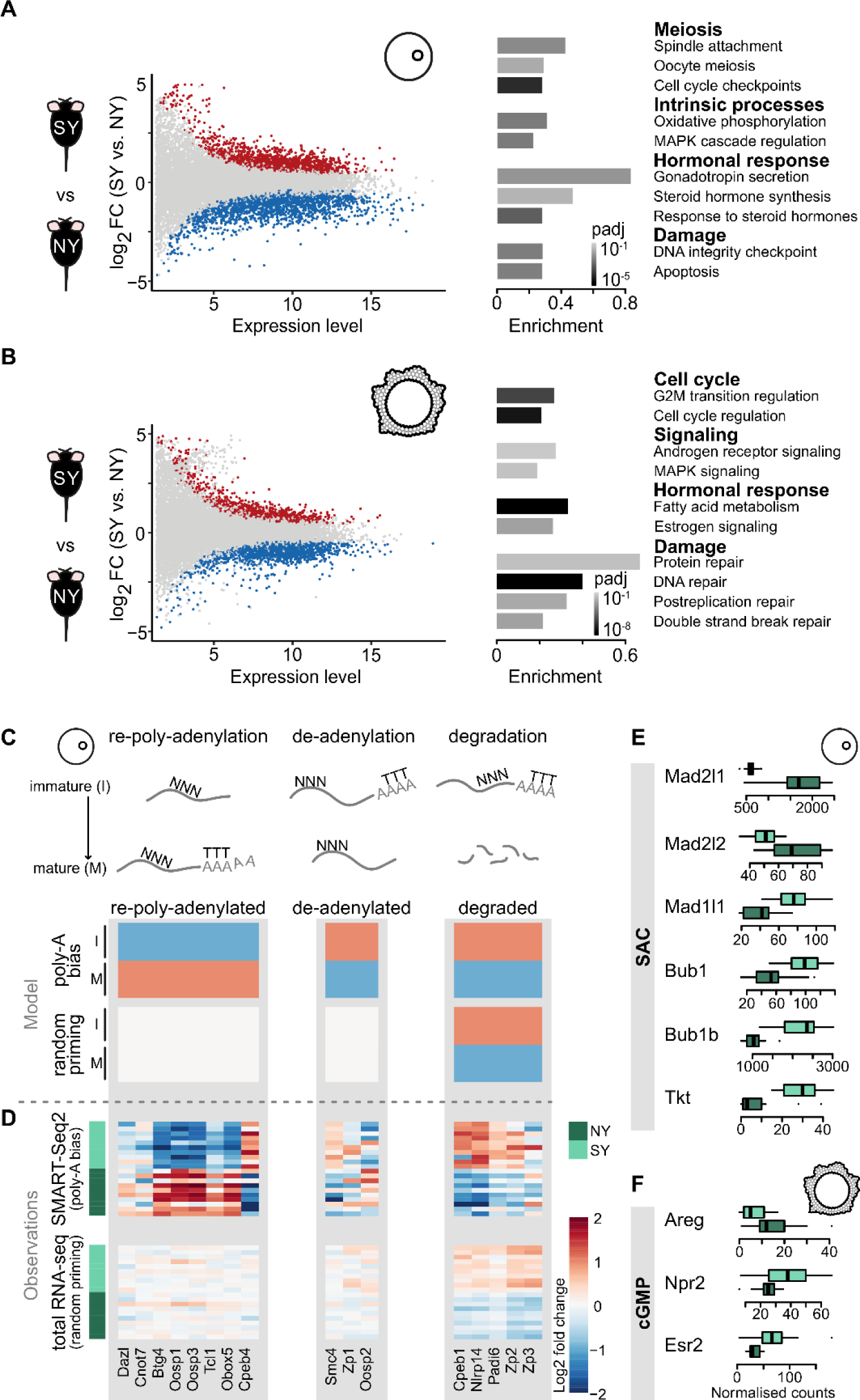
Superovulation disrupts the expression of genes involved in oocyte maturation. (A, B) Differentially expressed genes and enriched functional pathways between naturally ovulated and superovulated young oocytes (OCs) (A) and granulosa cells (GCs) (B). (C) Schematic representation of gene expression expectation in mature and immature oocytes between poly-A biased (SMART-seq2) and unbiased protocols (total RNA-seq) for genes involved in oocyte transcriptional remodeling during maturation (re-poly-adenylation, de-adenylation, and degradation). (D) Comparison of gene expression between naturally and superovulated oocytes using SMART-seq2 and total RNA-seq. For each technology, the fold change is computed between the two groups. Each row represents gene expression in an individual oocyte (randomly subsampled set). (E) Expression of genes involved in spindle assembly checkpoint (SAC) machinery in naturally and superovulated oocytes. (F) Expression of genes involved in the cGMP pathway in naturally and superovulated granulosa cells.

In addition to the unbiased analysis above, we compared the expression of genes involved in three mechanisms known to regulate oocyte function (cytoplasmic maturation, spindle assembly, and cell-cell communication) between natural and superovulated oocytes and their granulosa cells.

First, we inspected cytoplasmic maturation, and asked whether superovulation perturbs the expected re-poly-adenylation, de-adenylation, and degradation of known transcripts.^73–76^ To do so, we compared a poly-A-biased protocol (SMART-seq2) and a total RNA-seq protocol based on random priming of transcripts (Methods, Supplementary Table S5). An impairment of re-poly-adenylation of transcripts from superovulation would result in an apparent downregulation of these genes in the poly-adenylated biased experiments, whereas no difference would be observed between mature and immature oocytes in total transcriptome experiments (Figure 2C, re-poly-adenylation). Superovulated disruption of de-adenylation would create a symmetric pattern (Figure 2C). In contrast, degradation of transcripts should result in the same trend with both protocols (Figure 2C, degradation). Post-transcriptional regulators, including Dazl, Cnot7, Btg4, were previously identified as targets of re-poly-adenylation during oocyte maturation,^73,75–78^ and our experiments reveal that superovulation disrupts this process (Figure 2D). Similarly, we found that the de-adenylation of Smc4, a chromosome condensation gene whose poly-A tail is usually removed during oocyte maturation,^74^ is disrupted by superovulation. Finally, we found that the regulated degradation of transcripts during oocyte maturation is disrupted by superovulation, including the 3’-UTR binding protein Cpeb1.^73,76^ These results demonstrate that superovulation disrupts multiple interlinked mechanisms of transcriptome remodeling active in oocyte maturation.

Second, we asked whether superovulation perturbs the regulation of spindle assembly checkpoint (SAC) genes involved in meiosis. Genes involved in chromosome attachment to microtubules (Mad2l1, Mad2l2) were downregulated, while those responsible for accurate chromosome segregation (Bub1, Bub1b, and Ttk) were upregulated, which together suggest a delayed anaphase onset in superovulated oocytes (Figure 2E).

Third, we evaluated whether superovulation perturbs the cell-cell communication from granulosa cells controlling meiotic resumption in oocytes. In normal conditions, the luteinizing hormone (LH) surge upregulates *Areg* and downregulates *Esr2* and *Npr2* in granulosa cells. This inhibits cGMP production, leading to meiosis resumption in the oocyte.^79–82^ In our data, superovulation impaired this hormonal response of granulosa cells, likely affecting oocyte meiosis (Figure 2F).

In conclusion, superovulation perturbs germ and somatic cell transcription, affecting key COC maturation pathways.

### Superovulated COCs divide into two groups based on granulosa cell gene expression

We reasoned that the perturbations caused by superovulation would be first reflected within the extensive granulosa cell-oocyte communications which regulate oocyte maturation. For each COC, we constructed a cell-cell communication profile based on co-expression of ligand-receptor pairs (Methods). Naturally ovulated COCs grouped together based on hierarchical clustering, indicating homogenous cell-cell communication (Figure 3A, Supplementary Figure S2A). In contrast, we observed that superovulated COCs split into two distinct clusters (clusters S^N^ and S), where S^N^ more closely resembles naturally ovulated COCs. We further confirmed the relative similarity of S^N^ and N by dimensionality reduction analysis (Figure 3B).

**Figure 3.**
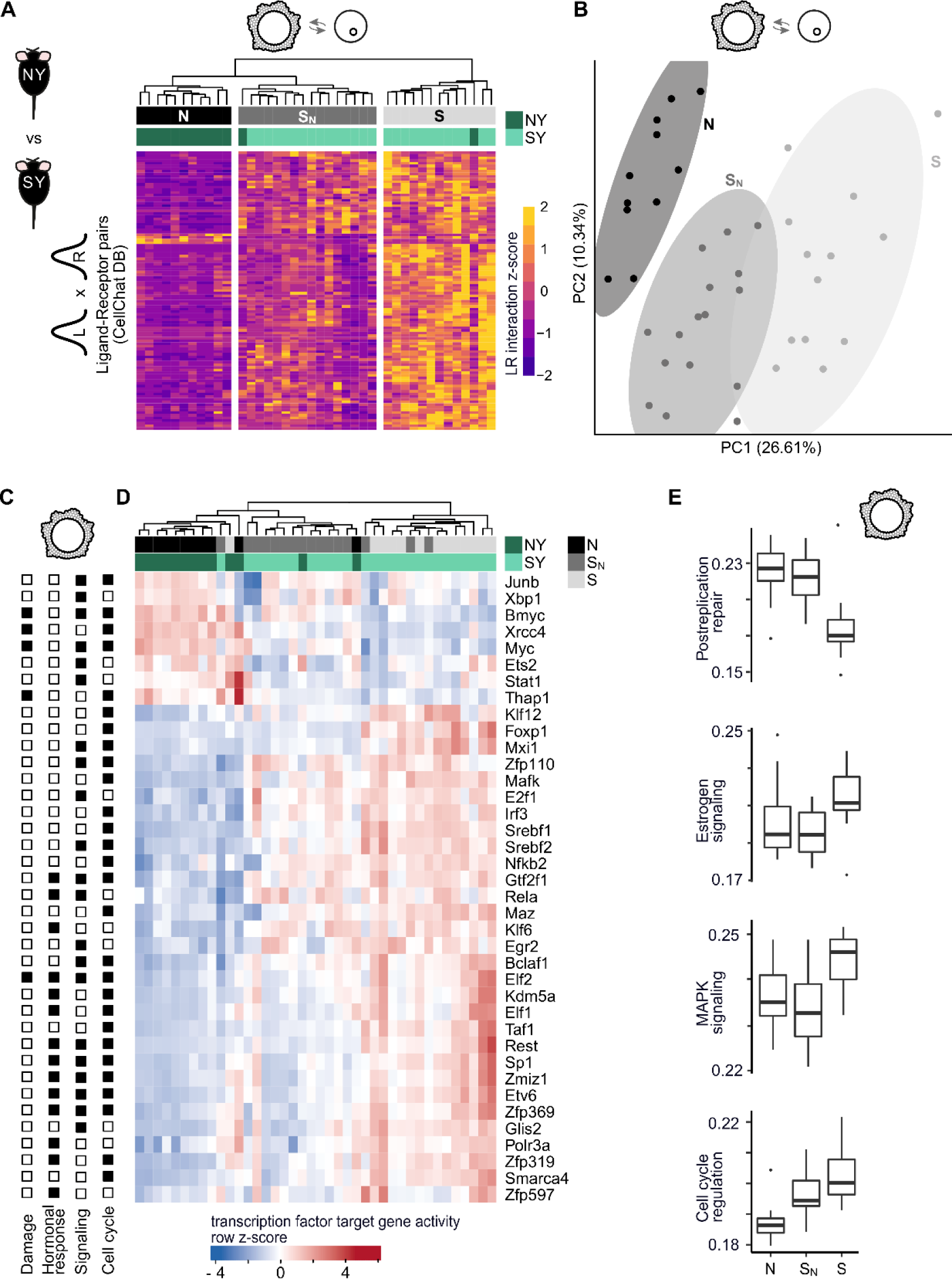
Superovulation dysregulates cell-cell communication and transcriptional regulation in cumulus-oocyte complexes. (A) Cell-cell communication scores between oocyte (OC) and granulosa cells (GC) calculated as a product of ligand and receptor gene expressions (Methods). Interaction scores are shown for ligand-receptor pairs (rows, z-scaled) in young natural and superovulated cells (columns) that displayed significant change between the three clusters (adjusted p-value < 0.05). Clustering dendrogram shown was obtained from the full list of ligand-receptor pairs shown in Supplementary Figure S3A. (B) PCA computed on OC-GC interaction scores. (C) Transcription factors associated with the pathways shown to be enriched in superovulated GCs in Figure 2B. Significant association with pathway categories (DNA damage, hormonal response, signaling, or cell cycle) is indicated as a filled square. (D) Activity scores of transcription factors whose activity scores were significantly different between naturally and superovulated young GCs are shown (permutation test, Methods). (E) Activity scores of selected pathways enriched in superovulated GCs from Figure 2B split by consensus clusters N, S_N_ and S. Activity scores for other pathways from Figure 2B are shown in Supplementary Figure S3D.

Next we reasoned that the similarity of S^N^ and normal COCs should be reflected in their corresponding transcription factor (TF) activities. First, we identified the transcription factors upstream of dysregulated COC pathways after superovulation, and calculated each TF’s activity score using the expression level of its target genes (Methods, Figures 3C and 3D). In oocytes, TF activity was grouped into natural and superovulated clusters without obvious substructure (Supplementary Figure S2B), as expected due to lack of active oocyte transcription at this meiotic stage (Su et al., 2007).

In contrast, superovulated granulosa cells again formed two distinct subclusters (clusters S^N^ and S, Figure 3D), paralleling the cell-cell communication results. These clusters were reproducible after bootstrapping and subsampling and were not the result of inter-individual variability (Supplementary Figure S2C). Pathway activity analysis further confirmed that S^N^ granulosa cells were highly similar to normally ovulated cells in replication and protein repair, MAPK and estrogen signaling, and had intermediate activity in cell cycle control, DNA repair and androgen receptor signaling (Figure 3E, Supplementary Figure S2D). Fatty acid metabolism and double strand break repair pathways were more similar between S^N^ and S compared to N. In all further analyses, we defined consensus clusters for N, S^N^, and S by including the 34 out of 40 granulosa cells where cell-cell communication and TF activity analyses were in agreement.

### Granulosa cell markers predict oocyte developmental outcome

We hypothesized that the two granulosa clusters may be linked to different embryo development outcomes. To link granulosa cell transcriptional state to early embryo development, we developed a targeted IVF assay (Methods). Briefly, we separated 67 new COCs into individual oocyte-granulosa cell pairs as previously and sequenced the granulosa cells. In parallel, we fertilized and cultured the corresponding single oocytes until early morula stage (86.6% fertilization rate), and assayed the transcriptome of the individual embryos (Figure 4A, Supplementary Figure S2E). Further, we trained a support-vector machine (SVM) classifier on the original 34 granulosa cells (Figure 3, 83.3 % accuracy on test set, Supplementary Figure S2F) and classified the granulosa cells from the new IVF experiment into S^N^, S and “unclassified” (Methods). We then transferred the labels from the classified granulosa cells to corresponding embryos and assessed embryos developmental progression, copy number variation (CNV), gene expression, and pseudotime.

**Figure 4.**
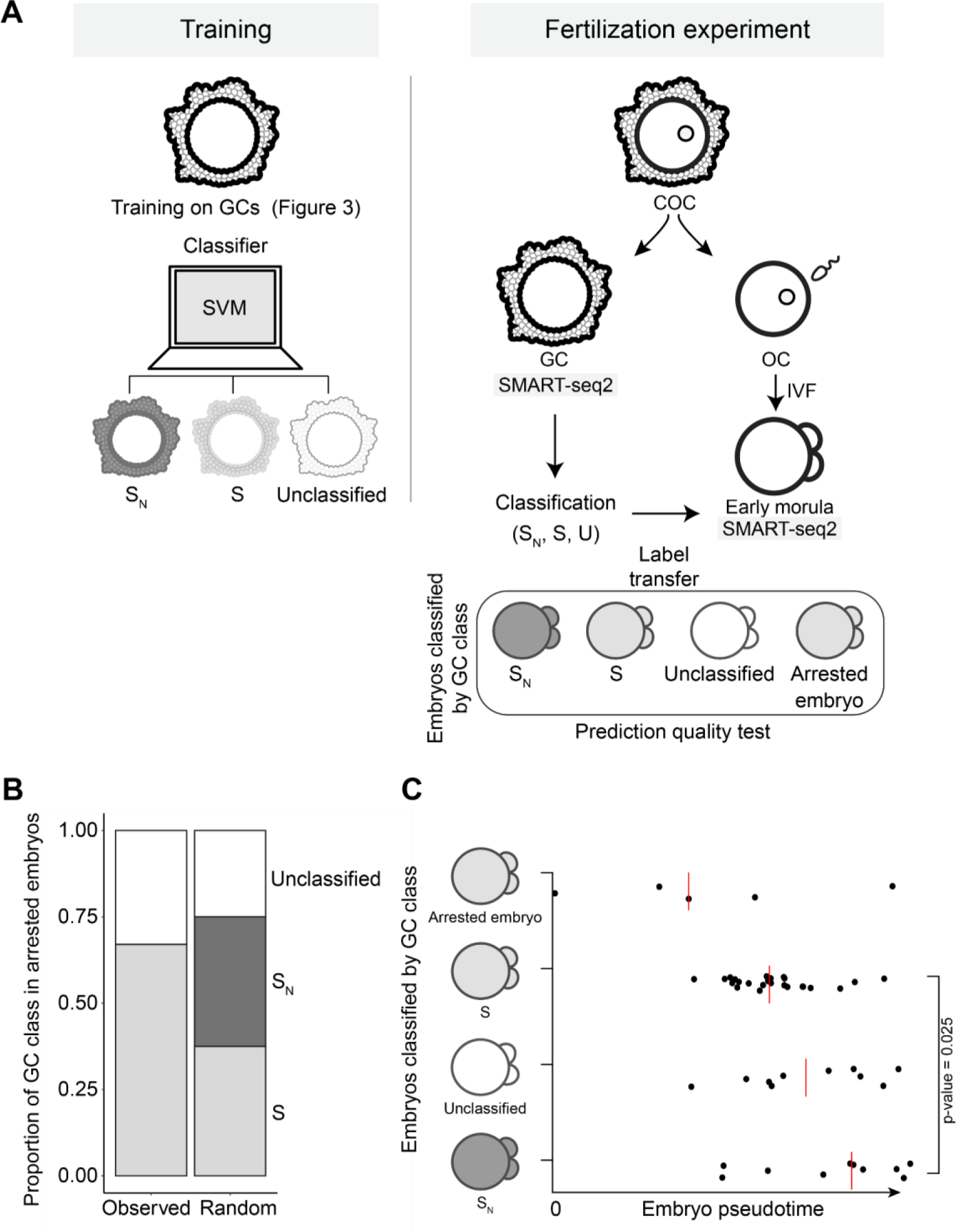
Pre-implantation embryo quality prediction from granulosa cells. (A) Schematic representation of targeted in vitro fertilization (IVF) experiment and embryo classification based on their granulosa cell transcriptomes. (B) Proportion of granulosa cell class in arrested embryos (n=6) in experiment and randomized sampling. (C) Embryos classified by their granulosa cell class along pseudotime calculated on highly variable genes. Red line -median. P-value = 0.025 between S_N_ and S classes, two-way Kolmogorov-Smirnov test.

Of these 58 embryos, 6 stopped developing before reaching the 8-cell stage (Supplementary Figure S2E), all of which were classified as S or “unclassified” (Figure 4B). Only one embryo had large enough aneuploidy detectable in RNA sequencing data (Supplementary Figure S2G) and it was classified as S. We then used pseudotime analysis of highly variable genes to compare embryos between classes (Methods). Compared to arrested embryos, the S^N^, S and unclassified embryos clearly separated (Figure 4C, S^N^ vs. S p-value = 0.025). Arrested embryos had the lowest pseudotime scores, followed by S and then S^N^.

In sum, the usefulness of our method is shown by its assignation of all embryos arresting before the morula stage to class S. Using this, we discovered that the original transcriptional state of a cumulus–oocyte complex is reflected in the gene expression of derived embryos, despite the massive genome remodeling occurring during fertilization and zygotic genome activation.^83,84^

### Aging and superovulation induce qualitatively similar transcriptional changes

Since both superovulation and aging lead to lower quality oocytes and poor developmental outcomes, we compared the mechanisms involved. Our experimental design newly permitted a rigorous comparison of how COC transcription changes during natural aging (that is, naturally ovulated young versus old cells) and upon superovulation (that is, naturally ovulated young cells versus superovulated cells) (Figure 5A). We discovered that in oocytes and granulosa cells both superovulation and aging perturb a common set of genes and with a similar directionality (Figures 5B and 5C, Supplementary Figure S3A and S3B).

**Figure 5.**
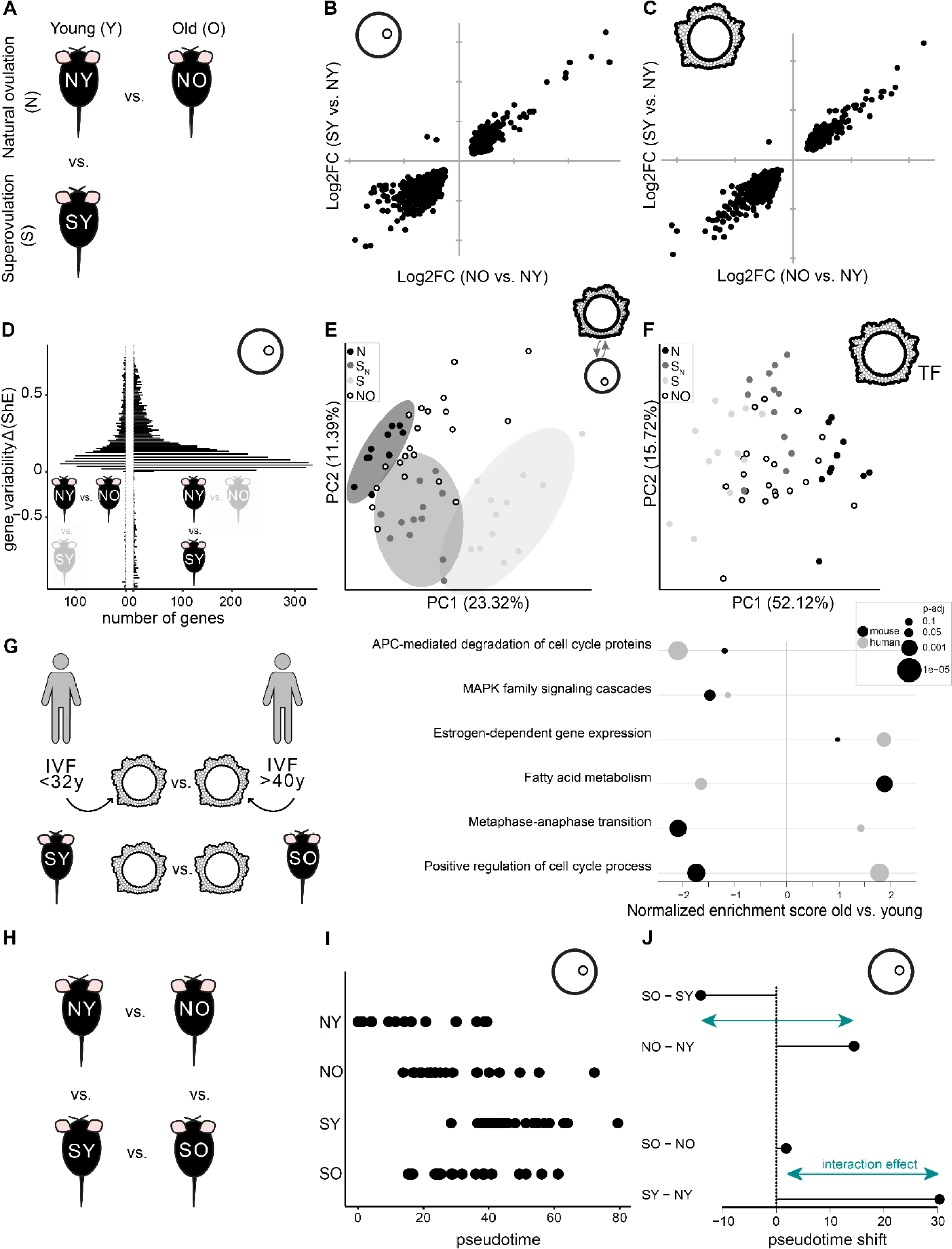
Superovulation and aging lead to similar but non-additive transcriptional changes in cumulus-oocyte complexes. (A) Schematic representation of the natural aging and superovulation comparison. (B, C) Log2 fold change of gene expression in SY versus NY compared to Log2 fold change in NO versus NY oocytes (B) and granulosa cells (C). Only genes that were significantly differentially expressed in both comparisons are shown (adjusted p-value < 0.05), all genes are shown in Supplementary Figure S3A and S3B. (D) Distribution of gene variability values calculated as differential Shannon entropy in NO (left) and SY (right) oocytes in comparison to NY oocytes. Increased Shannon entropy is associated with increased gene expression variability. (E) PCA computed on ligand-receptor interaction scores between NY, SY, and NO oocytes and granulosa cells (clusters N, S^N^ and S defined in Figure 3). (F) PCA computed on the activity scores of the transcription factors shown in Figure 3D in NY, SY, and NO granulosa cells (clusters N, S^N^ and S defined in Figure 3). (G) Selected dysregulated pathways in granulosa cells during aging; young - 3M for mice (n = 4), <32y for human patients (n = 4) and old - 12M for mice (n = 3), >40y for patients (n = 4), full list of pathways in Supplementary Table S6. (H) Schematic representation of the 4-way comparison of aging and superovulation. (I) Pseudotime analysis computed on highly variable genes in mouse oocytes in all four experimental conditions. (J) Quantification of shifts in pseudotime between groups contrasted in Figure 5I. For each group the mean pseudotime was used to compute the shifts. The interaction effect represents the non-additivity of aging and superovulation.

We asked whether aging may also increase cell-to-cell variability in oocytes, as has been previously reported for other cell types.^85–88^ We first compared gene expression variability by Shannon entropy between young and old oocytes (Figure 5D), and found that aging increases cell-to-cell variability. Similar analyses revealed that superovulation also substantially increases oocyte cell-cell variability in gene expression (Figure 5D).

To confirm that gene expression variability is reflected in upstream and downstream communication and TF usages, we then computed cell-cell communication and TF activity profiles for old COCs. We observed that old COCs locate between young natural and superovulated COCs in communication profile space (Figure 5E). Likewise, TF activity profiles of old granulosa cells localized close to cluster S^N^ in TF activity space (Figure 5F).

Together, these results strongly suggest that superovulation and aging lead to quantitatively but not qualitatively different responses, targeting the same underlying regulatory networks.

To investigate whether similar aging effects can be observed in human, we compared the pathways enriched during aging in superovulated granulosa cells from mice and patients undergoing IVF (Figure 5G). Despite possible differences arising from the cell collection protocol (in-ovary granulosa cells in human and ovulated granulosa cells in mouse), the dysregulation of signaling and response to hormone pathways was conserved, suggesting that aging might similarly impact COC quality in humans. In contrast, dysregulation of cell cycle and fatty acid metabolism diverged, among others (Figure 5G, Methods, Supplementary Table 6).

### Superovulation and aging effects are non-additive

Finally, we investigated whether the effects of superovulation and aging on COCs are cumulative (Figure 5H). Surprisingly, among old oocytes and granulosa cells, we found very few significant differences in gene expression levels between naturally and superovulated cells (Supplementary Figures S3C and S3D). This analysis had sufficient statistical power to identify any gene expression differences, as the number of replicates was comparable to the young mouse experiments (Supplementary Table 1). Thus, aging and superovulation do not have an additive effect on gene expression. By ordering the cells based on highly variable genes in a pseudotime analysis, we observed that the impact of superovulation is reduced in old COCs (Figure 5I, Supplementary Figure S3E). We reasoned that this would also be reflected in oocyte maturation. We computed gene expression of maternal remodeling targets, cGMP cascade genes and SAC genes in old oocytes or granulosa cells (Supplementary Figures S3F and S3G) and indeed found that old COCs often exhibit intermediate gene expression profiles.

To test the relative impact of age and superovulation, we quantified the shifts in pseudotime between all four groups (NO, SO, NY, SY) (Figure 5J). We found that superovulation induces a strong transcriptional shift in young COCs, which is very attenuated in the old cells. Meanwhile, age-induced transcriptional changes are different in natural and superovulated COCs, indicating that experiments performed with superovulated cells are not representative of aging in naturally ovulating mice.

## Discussion

Our study introduces two important technical refinements that allowed us to quantitatively dissect the impact of superovulation and aging on COCs in mice. First, we report a novel experimental approach to analyze the functional interactions between oocytes and their surrounding granulosa cells, based on careful manual isolation. Second, our experimental design incorporates COCs from naturally ovulated older mice, introducing a key control missing from most studies to date.

The efficiency of fertility treatments is limited by the routine retrieval of a large number of immature oocytes that cannot be fertilized. *In vitro* maturation of these oocytes is a promising solution to obtain more embryos without increasing the number of ovarian stimulation cycles.^89^ Understanding where the oocyte-granulosa cell communication and oocyte maturation cues are lost is critical to efficiently design such protocols.

Our transcriptomic approach allowed us to confirm previously predicted meiotic perturbation of oocytes in superovulated COCs.^37,90–93^ For the first time, we show that maternal transcriptome remodeling is also impaired in superovulated oocytes. This could explain the delayed development observed in resulting embryos^39,40,94^ and cause embryo developmental arrest if zygotic genome activation is prevented.^95–98^ We suggest that insufficient maturation of the oocytes is caused by aberrant granulosa-to-oocyte communication through the disruption of the cGMP signal transduction pathway - an essential pathway for meiosis resumption.^99^ We hypothesize that granulosa cell failure to respond to and transmit ovulation cues to the oocyte is responsible for poor oocyte quality. More work is still needed to understand whether granulosa cell response to LH is simply delayed or if they are unable to properly relay maturation signals.

Granulosa cell transcription has been studied as a possible non-destructive readout to identify the best quality oocytes in IVF. However, prior attempts to identify gene expression signatures in granulosa cells have been retrospective where embryo developmental outcome was used to assess the quality of granulosa cells. Our design enabled us to undertake a prospective study, where we used granulosa cell transcriptomes to group the embryos and study their development. Moreover, our inclusion of a naturally ovulated group enabled us to separate superovulated granulosa cells into two groups. Our granulosa cell classifier was able to successfully predict a population of embryos with a low pseudotime that included all arrested and aneuploid embryos.

Aging is also associated with a lower oocyte quality.^100^ Prior analyses of age impact on oocytes have only ever been assessed in superovulated cells.^7,8,10^ By including naturally ovulated cells, we were able to decouple the impact of aging and superovulation. Our systematic comparison revealed that aging and superovulation induce similar transcriptional perturbations in oocytes and granulosa cells. One explanation for this similarity is that exogenous gonadotropin stimulation in young individuals may mimic age-related increase of gonadotropin levels in both humans and mice.^101–103^ Despite their similarity, hormone and age effects are non-additive, implying that superovulation and aging are two processes that do not add up linearly, but influence each other. Superovulation has a greater transcriptional impact in younger COCs, possibly reflecting reduced sensitivity of granulosa cells to gonadotropins in old age.^104^ These data are also relevant for human IVF patients as we show that aging induces similar transcriptional changes in signalling and hormonal response pathways in their granulosa cells.

Importantly, any studies attempting to identify the impact of aging on oocyte fertility must take into account the non-additive effects of aging and superovulation. In other words, the effects of natural aging cannot be inferred only by analyzing superovulated oocytes, and the inclusion of experiments using naturally ovulated older female mice is highly recommended.

Our study thus reveals how the transcriptional state of a COC both shapes and survives the oocyte-to-embryo transition, and that aging and superovulation dysregulate the same pathways to degrade fertility.

## Methods

### Animals

Naturally ovulated or superovulated oocytes and granulosa cells were collected from 11-14 weeks (young) and 50-58 weeks (old) *Mus musculus* (C57BL/6J and C57BL/6Ly5.1) female mice. Each experimental group contained 3-4 mice. Mice were purchased from Janvier (France) and allowed to adapt to animal facility conditions for at least 1 week. Mice were kept at the DKFZ animal facility under controlled light-dark cycle (12h/12h, from 7:00 to 19:00) and had access to standard laboratory chow and water ad libitum. Around half of old mice presented female reproductive tract pathologies and therefore could not ovulate. All animal experiments were carried out according to governmental and institutional guidelines and approved internally (DKFZ-366) and by the local authorities (Regierungspräsidium Karlsruhe, G-238/19).

### Induction of ovulation

Ovulation by hormonal stimulation was induced by intraperitoneal injection of 5 IU pregnant mare serum gonadotropin (PMSG - Pregmagon®) and a second injection of 5 IU human chorionic gonadotropin (hCG - Ovogest®) 48 hours later.^105^ The COCs were collected from the oviduct ampullas 14-16h after hCG injection in young mice and 18-19h in old mice. Superovulated young mice yielded 13 healthy oocytes on average, whereas old mice yielded 11 (Supplementary Table S7). Oocytes were considered healthy if they had a spherical shape with a uniform translucent cytoplasm and were surrounded by expanded granulosa cells.

Natural ovulation was induced by mating C57BL/6 females with vasectomized CD-1 males.^37,106,107^ The mice were caged in a 1:1 female-male ratio shortly before the beginning of the dark cycle. The next morning females presenting a copulatory plug were sacrificed for cell collection. Young mice were sacrificed 2-3h after the beginning of the light cycle, and old mice after around 6h. Females without plugs were examined each morning for three consecutive days. Males were allowed to rest for at least one day between matings. Naturally ovulated young mice yielded 7 healthy oocytes on average, whereas old mice yielded 9 (Supplementary Table S7).

### COC extraction

Mice were sacrificed by cervical dislocation and the oviducts dissected. Under the stereomicroscope (SteREO Discovery.V12, Zeiss) ampullas were torn to release the COCs into a 96 µl M2 media (M7167, Sigma) drop under mineral oil (69794, Sigma) at room temperature. Then, 4 µl of pre-heated 500 µg/ml hyaluronidase (H3884, Sigma) diluted in M2 media (final concentration in the drop 20 µg/ml) was added to separate the COCs into single units. The COCs were incubated for 10-20 min at 37°C and then mechanically separated into individual M2 drops using a 115-124 μm glass retransfer pipette (BioMedical Instruments). Individual COCs were then washed in M2 once and incubated for less than 5 min with enzymatic mix Accutase (A6964, Sigma) (Zhang et al., 2018) at 37°C to further separate the granulosa cells from the oocytes. Once separated, GCs (∼50-200) were collected from these individual drops as a small bulk using a mouth pipette pre-filled with Dulbecco’s phosphate-buffered saline (DPBS, 59321C, Sigma). Oocytes were washed twice with M2 media and once with DPBS before collection. Cells were immediately flash frozen in liquid nitrogen in individual 0.2ml thin wall PCR tubes (731-0679, VWR) and stored at -80°C until further use (Supplementary Figure S1). Only oocytes with uniform translucent cytoplasm and spherical shape were collected. The media in the mouth pipette was changed between micro-manipulations of singularized COCs, granulosa cells and oocytes to ensure sample specificity. All samples were collected within 1.5 hours after mouse dissection.

### SMART-seq2

Oocyte and granulosa cell samples were processed for full-length cytoplasmic RNA amplification using a slightly modified SMART-seq2 protocol.^108^ Libraries for next generation sequencing were prepared using Nextera XT DNA library Prep Kit (Illumina). Briefly, cells were lysed with 0.2% Triton X-100 (Sigma Aldrich) and reverse transcription was facilitated by Maxima-H Minus Reverse Transcriptase (Thermo Fisher). Amplification of complementary deoxyribonucleic acid (cDNA) was achieved with 17 PCR cycles for oocytes and 20 cycles for granulosa cells. 1 ng of cDNA was used as template for library preparation, then tagmented, double indexed and amplified with 11 PCR cycles. All libraries were multiplexed into a single pool prior to sequencing. Quality control for sample preparation was performed using 4200 TapeStation High Sensitivity DNA and D1000 tapes (Agilent), and Qubit 4 fluorometer High Sensitivity dsDNA assays (Thermo Fisher). Samples that failed cDNA amplification were excluded. Samples were sequenced on NextSeq 550 (Illumina) using Single-Ended 75bp High-Output kits at the DKFZ Open Sequencing Lab.

### Targeted *in vitro* fertilization

Young C57BL6/J mice were superovulated and COCs singularized as described above. Granulosa cells were immediately flash frozen and paired oocytes were divided into individual drops of CARD media (Cosmobiousa) under mineral oil (Vitrolife) for fertilization. The oocytes were incubated at 37°C, 5% CO^2^ for 30-40 min prior to fertilization.^109^

Frozen *Mus musculus* (C57BL6/J) male sperm was obtained from the DKFZ Transgenic Service. The sperm was frozen as described previously^110^ where sperm straws contained 6.41-10.6 x 10^6/ml progressive spermatozoids with 7.75-14 x 10^6/ml motility. The sperm straws were thawed and capacitated in Fertiup media (Cosmobiousa) for 30 min as recommended by the manufacturer.

Fertilization was achieved by combining 2.5 µl of activated sperm with 20 µl CARD media drops containing a single oocyte and incubated at 37°C, 5% CO^2^ for 3 hours. Oocytes were then individually washed in Human Tubal Fluid (HTF) media drops and transferred to 20 µl HFT drops for overnight culture. After 24 hours, the fertilization rate was evaluated. 2-cell stage embryos were then transferred to G1+ media (Vitrolife) for further development. Early morula stage embryos were continuously collected based on an observation window of 62 to 64h post-fertilization (Supplementary Figure S3). For collection, the embryos were washed in DPBS and flash frozen as in the method section “COC collection”. Granulosa cells and embryo samples were further processed as described in method section “Smart-seq2”, except that cDNA amplification PCR cycle number for embryos was 16-17. All samples were sequenced on a Novaseq 6000 at the Institute of Clinical Molecular Biology (IKMB), Kiel, Germany.

### Total RNA sequencing of oocytes

Oocytes from naturally and superovulated mice were collected as described in the sections “Animals” and “Induction of ovulation”. COCs were singularized as described in the methods section “COC extraction”. SMARTer Stranded Total RNA - Low Input Mammalian (Takara) kit was used for lysis, cDNA amplification and library preparation with 10 cycles for PCR-1 and 16 cycles for PCR-2. Samples were sequenced as recommended by the manufacturer on Novaseq 6000 with paired-end sequencing at IKMB.

RNA processing from human granulosa cells

Ovarian stimulation was performed depending on the individual patient’s situation as previously described.^111^ Granulosa cells were retrieved from the follicular fluid after transvaginal ultrasound-guided follicle puncture for IVF as previously described.^111,112^ Patient procedures were carried out at the Universitäts-Frauenklinik, Heidelberg, Germany. All participating patients signed informed written consent and completed clinical questionnaires. All patient procedures were approved by the local ethical committee of the University of Heidelberg, Germany (S-602/2013).

Total RNA was extracted from 4 young (⩽32 years) and 4 older patient (⩾40 years) granulosa cell samples using TRIzol (Life Technologies, Carlsbad, CA, USA) as previously described.^111,112^ Further, RNA was cleaned from DNA with DNA-free™ DNA Removal Kit (Invitrogen). ∼1 µg of RNA was used to construct libraries with Truseq Stranded mRNA kit (Illumina) and barcoded with IDT for Illumina-TruSeq DNA and RNA UD Indexes. Samples were sequenced on Nextseq 2000 (Illumina) with 100bp paired-end sequencing at DKFZ, Heidelberg, Germany.

### Sequencing quality control

Sequencing raw alignment files were demultiplexed by the DKFZ Genomics and Proteomics Core Facility. The adaptors were then trimmed and the sequencing reads were aligned to the mm10 genome (GRCm38) using STAR (v.2.7.0f). Gene-barcode count matrices were then analyzed using R (v.4.0.0) and Seurat^113^ (v.4.0.3).

Samples were kept if the number of reads was above 5000, the number of genes detected was above 1000, and the percentage of mitochondrial reads was below 5% (Supplementary Table S1). Cell-type-specific marker gene expression was checked and samples that were mislabeled were removed (10 samples in total) and the corresponding libraries resequenced.

Raw count tables for all replicates were merged and normalized together using the SCTransform function from Seurat with default parameters. PCA was computed using top 3000 most variable genes and UMAP was computed on the first 30 PCs.

### Differential gene expression (DGE) analysis

DGE analysis was performed using DESeq2^114^ (v.1.28.1) and a pseudo-bulk approach. Briefly, SCT-corrected counts of each mouse were summed across all cells from the same cell-type and used as the input for DESeq2. The effects of age and superovulation were modeled with an interaction term between age and superovulation.

### Overrepresentation analysis (ORA)

ORA was performed using a hypergeometric test. Gene sets used were downloaded from the MsigDB (https://www.gsea-msigdb.org/gsea/msigdb/) pathway collection (Hallmarks, Kegg, Reactome and GO Biological Process). P-values were adjusted using the Benjamini-Hochberg procedure.

### Cell-cell communication (CCC) analysis

To assess the CCC between OC and GC within each follicle we computed an interaction score for each pair based on the CellChat^115^ ligand-receptor database and gene expression. Gene counts were corrected using the SCnorm method (v. 1.10.0) to take into account gene length. Only the top 100 most expressed ligand and receptor genes were used for analysis. Interaction scores were computed by multiplying the expression of the ligand gene in the sender cell and of the receptor gene in the receiver cell. In case of receptors with multiple subunits, only the least expressed subunit was considered. For each OC-GC pair, communication was assessed between OC and GC in both directions, as well as GC-to-GC and OC-to-OC (autocrine signaling).

### Single-Cell Regulatory Network Inference and Clustering (SCENIC) analysis

SCENIC^116^ (v. 1.2.4) was used to perform single-cell regulatory network analysis in granulosa cells using the SCT normalized gene expression values. Gene co-expression networks were determined using GENIE3, enriched transcription factor motifs were scored using RcisTarget and regulon activity scores were calculated using AUCell. To assess which regulons overlap with the pathways found to be significantly dysregulated by superovulation, we used a Jaccard index to quantify the overlap between a regulons’ target genes and the genes in each pathway. A hypergeometric test was used to assess the enrichment. Hierarchical clustering of transcription factors was performed using Euclidean distances and complete clustering method.

### Pathway activity scoring

Scoring of pathway activity was performed using the AUCell package^116^ (v. 1.10.0) by calculating an enrichment score of each gene set within the top 20% most expressed genes in each cell. The gene sets used in this analysis were the same as gene sets used in the ORA.

### Differential Shannon’s entropy

Differential Shannon Entropy (dShE) was used to assess the differences in transcriptional heterogeneity between young and old naturally ovulated oocytes as well as superovulated and naturally ovulated young oocytes. Differential ShE was calculated using the EntropyExplorer package^117^ (v. 1.1). Multiple testing correction was performed using the Benjamini-Hochberg procedure.

### Pseudotime analysis

Slingshot^118^ (v.1.6.1) was used to calculate pseudotime trajectories along aging, in superovulation, as well as for embryos. Principal component analysis was used to perform dimensionality reduction. Cells were clustered using Gaussian mixture modeling (package mclust, v. 5.4.10), and the principal curves fitted through cell clusters using the Slingshot function.

### Total RNA-seq analysis

Sequencing raw alignment files were demultiplexed by the DKFZ Genomics and Proteomics Core Facility. The sequencing reads were aligned to the mm10 genome using the Cogent NGS Analysis Pipeline (CogentAP, Takara Bio, v.1.5.1). Gene-barcode count matrices were then analyzed using R (v.4.0.0) and Seurat (v.4.0.3).

Samples passed a quality filter if the number of genes detected was above 5000 and the percentage of ribosomal and mitochondrial reads was below 18 and 2.5%, respectively. Cell-type-specific marker gene expression was checked to exclude granulosa cell contamination.

Raw counts for all replicates were merged and corrected using the SCTransform function from Seurat. Genes of interest targeted by maternal transcriptome remodeling were identified based on previous literature. For each sample and each gene we computed a log2 fold change using the average expression of the gene in the other ovulation group (natural ovulation was used as a reference for superovulated samples, superovulation as a reference for naturally ovulated ones).

### IVF analysis

We trained two support vector machines to classify the new granulosa cells from the IVF experiment into S^N^ and S groups. We used the consensus S^N^ and S clusters of granulosa cells from the original experiment as training and test set. The package caret (v. 6.0-94) was used to perform the classification using support vector machines with a linear kernel.

For the first classifier, we used as input training set the transcription factor activity scores calculated using SCENIC as detailed above. Only transcription factors that were determined to regulate the pathways perturbed in superovulation (see Single-Cell Regulatory Network Inference and Clustering (SCENIC) analysis) were considered in the analysis. To further refine the selection of transcription factors we calculated which factors showed significantly differential activity scores between the S^N^ and S groups using a generalized linear mixed model with random intercept.

Activity scores were used as the dependent variable, while group labels were used as the independent variable, and sample label as random effect. The model was fit under a beta distribution using the glmmTMB package (v. 1.1.7). P-values were corrected for multiple testing using the Benjamini-Hochberg procedure. Transcription factors that showed significantly different activity (adjusted p-value < 0.05) between the two groups were used as features to train the classifier.

For the second classifier, we used DESeq2 to identify differentially expressed genes between the S^N^ and S groups, while taking into account mouse of origin. Differentially expressed genes that passed the threshold (baseMean above 50, adjusted p-value below 0.01 and absolute log2 fold change above 0.7) were used to compute a PCA. PC 1 clearly separated the S^N^ and S groups and was thus used to further filter the genes (loading above 0.09, roughly top 50% genes). Genes that passed these thresholds were used to train the second support vector machine classifier. Prior to training, features that showed near-zero variance, that were correlated or showed linear dependency were removed. A genetic algorithm was used to perform the final feature selection. To tune the cost hyperparameter we used adaptive resampling.

Superovulated granulosa cells from the IVF experiment were then classified using both the transcription factor and the differential gene expression trained classifiers. If the classification label of the IVF granulosa cells was different between the two classifiers then the cells were labeled as unclassified.

Embryos which developed from matching fertilized oocytes were split in two groups based on their granulosa cell classification. The InferCNV package (v. 1.17.0) was used to detect copy number variations in the embryos. Embryos classified as S^N^ were used as a control group.

To calculate differential expressed genes between S^N^ and S embryos we fitted a negative binomial generalized linear model using the MASS package. Multiple testing correction was performed using the Benjamini-Hochberg procedure.

### Human data analysis

Sequencing raw alignment files were demultiplexed by the DKFZ Genomics and Proteomics Core Facility. The sequencing reads were aligned by the DKFZ Omics IT and Data Management Core Facility (ODCF) to the 1KGRef_PhiX genome (hs37d5 genome from the 1000 genomes project with the PhiX-sequence as an additional contig) using STAR (v.2.5.3a) and the DKFZ ODCF RNAseqWorkflow pipeline (v.1.3.0). Gene-barcode count matrices were then analyzed using R (v.4.0.0).

Samples were filtered based on mitochondrial reads percentage (below 18%, corresponding to 80% percentile) to remove low quality samples.

DGE analysis was performed using DESeq2 (v.1.28.1) and age group as a predictor (below 32 years old or above 40).

Gene set enrichment analysis was performed using the fgsea package^119^ (v.1.14.0) using the same pathways as before (Hallmarks, Reactome and GO Biological Process, see Methods Overrepresentation analysis) and log2 fold change from DESeq2. The same analysis was performed on the mouse SO vs SY DGE results as a comparison. All pathways that are significant (adjusted p-value < 0.05) in at least one species are available in supplementary.

## Data availability

Data is available in ArrayExpress under accession numbers E-MTAB-13479 (paired oocytes and granulosa cells SMART-seq2), E-MTAB-13480 (paired morulas and granulosa cells SMART-seq2), E-MTAB-13474 (total RNA-seq), and E-MTAB-13496 (human data).

## Code availability

Code used in this study is available in github repository: https://github.com/goncalves-lab/follicle_project.

## Acknowledgements and Funding

We thank Open Sequencing Lab (DKFZ), Institute of Clinical Molecular Biology (IKMB), U. Bender (Universitäts-Frauenklinik Heidelberg), I. C. R. Delgado (EMBL), animal caretakers from ATV-108, U. Kloz, L. Ziegler, F. van der Hoeven and R. Brecht (DKFZ) for technical support and assistance. We thank P. A. Ginno for offering analysis suggestions. This work was supported by NCT/Helmholtz core funding (B270 to D.T.O.and B210 to A.G.); European Research Council (788937 to D.T.O.); Helmholtz junior group leader post (to A.G.); and the Deutsche Forschungsgemeinschaft (FI 2558/1-1 to A.G. and RE 3647/1-2 to J.R.).

## Author contributions

Conceptualization, K.D., P.L., A.G., D.T.O.; Methodology, K.D., P.L., I.W.; Investigation, K.D., P.L., I.W., N.S., A.S., M-L.K.; Formal Analysis, P.L., I.W.; Resources, A.T., X.P.N., A.V., J.R.; Data Curation, K.D., P.L., I.W.; Writing – Original Draft, K.D., P.L., I.W., A.G., D.T.O.; Visualization, K.D., P.L., I.W.; Project Administration, K.D.; Funding Acquisition, A.G., D.T.O., J.R.; Supervision, A.G., D.T.O. All authors had the opportunity to edit the manuscript, and all authors approved the final manuscript.

## Materials & Correspondence

Correspondence and materials requests should be addressed to Duncan T. Odom and Angela Goncalves.

