## Supplementary Figures for "Superovulation and aging perturb oocyte-granulosa cell communication"

### Supplementary data

#### Supplementary figures

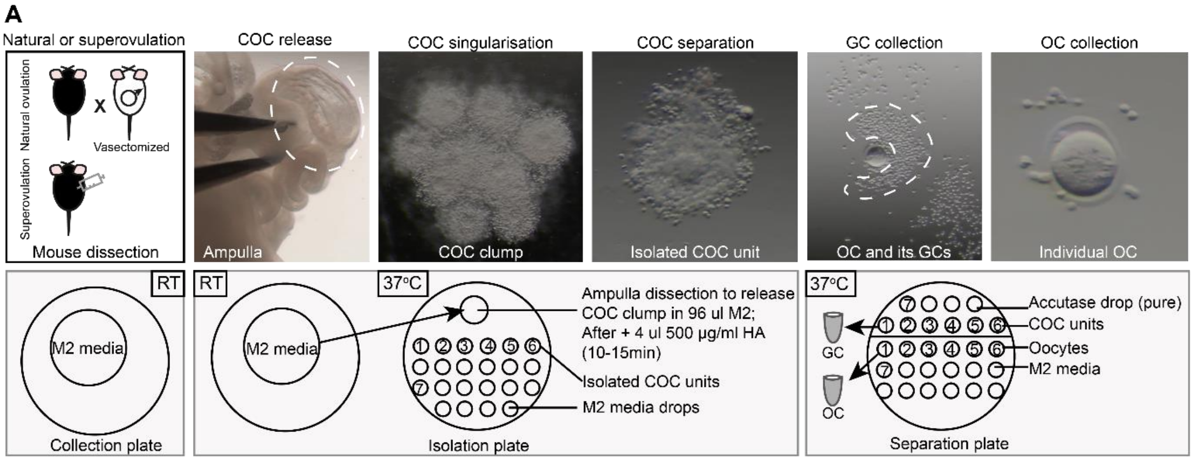


**Figure S1.** (A) Experimental protocol for COC singularization and paired oocyte-granulosa cell separation.

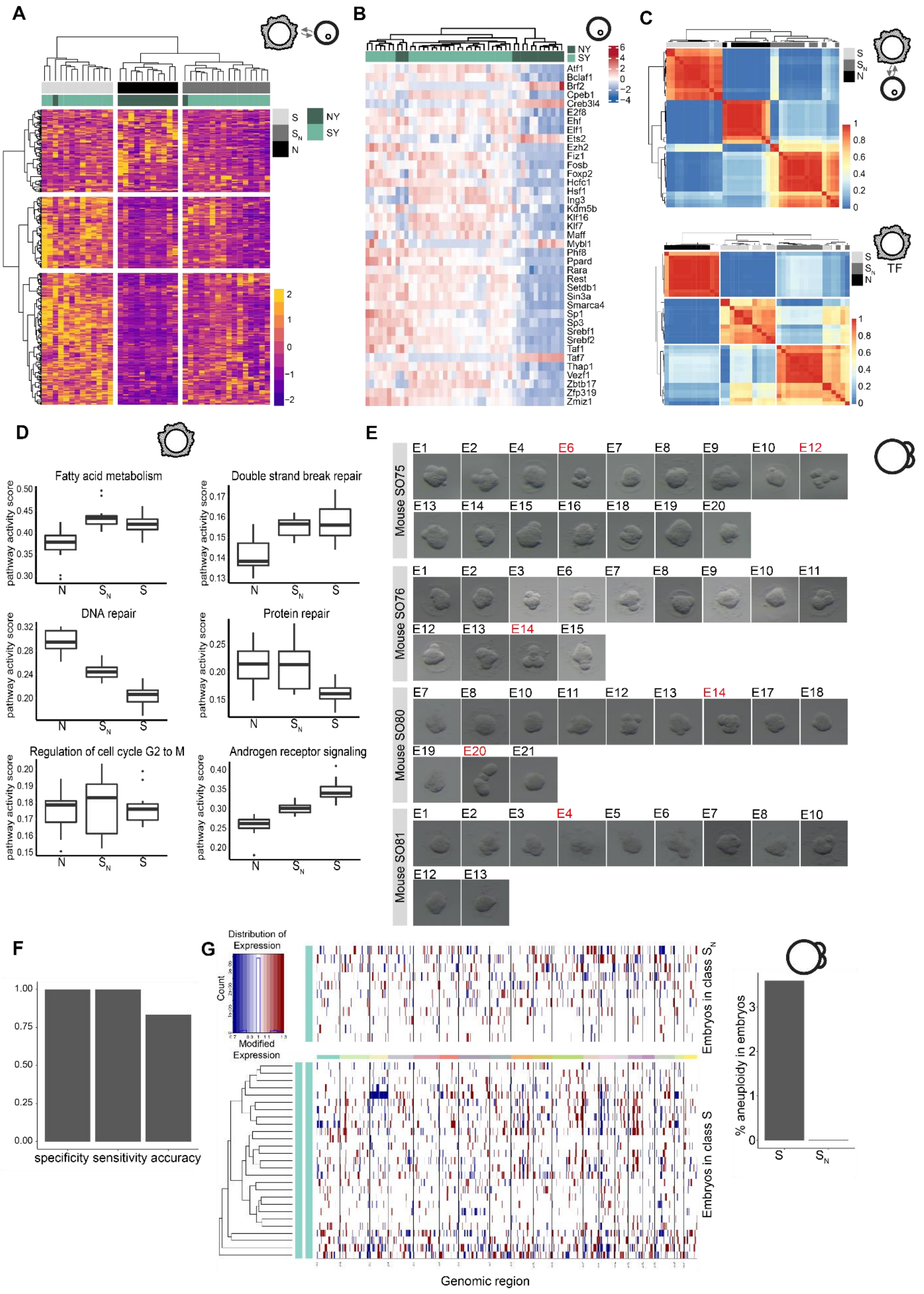




**Figure S2.** (A) Cell-cell communication scores for all ligand-receptor interactions tested. Each row represents a ligand-receptor interaction and each column is a granulosa-oocyte pair. (B) Activity scores of transcription factors associated with enriched pathways shown in Figure 2A in OCs. TFs whose activity scores were significantly different between naturally ovulated and superovulated young OCs by a permutation test are shown. (C) Stability of the clusters obtained from cell-cell communication (top) and transcription factor (bottom) analyses assessed using bootstrapping. The color scale represents the frequency of co-clustering of two samples. (D) Activity scores of enriched pathways from Figure 2B in GCs split by consensus clusters N, $S_N$ and S identified in Figures 3A and 3D. (E) Widefield images of all embryos collected at the morula stage. Embryos which arrested before the morula stage are labeled in red. (F) Specificity, sensitivity and accuracy of the GC classifier on the test set. (G) Whole genome copy number representation from inferCNV using SN embryos as a reference (Methods). One embryo was identified as aneuploid and was classified as S based on its granulosa cells transcriptome.

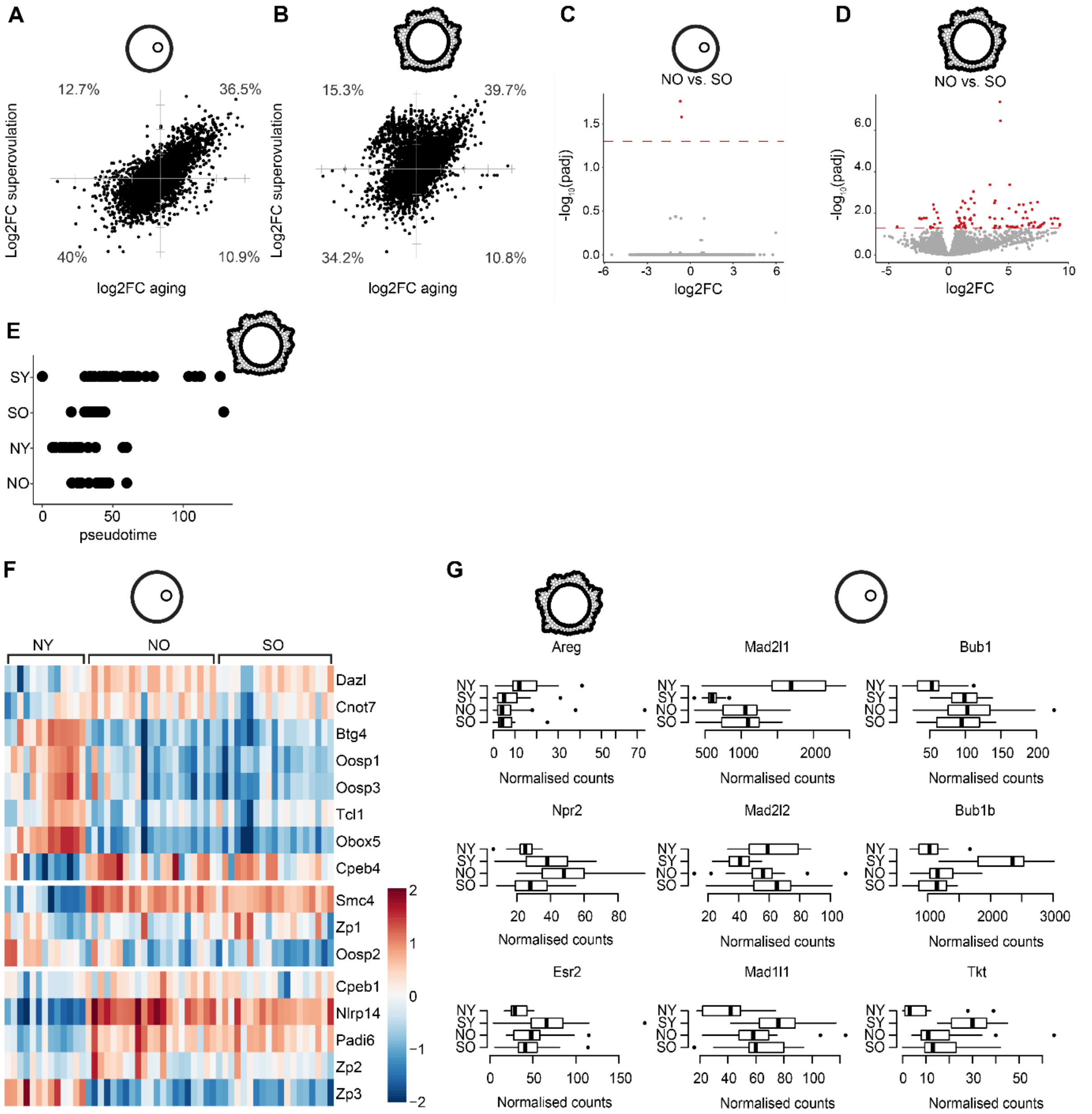


**Figure S3.** (A, B) Log2 fold change of gene expression in SY versus NY compared to log2 fold change in NO versus NY oocytes (A) and granulosa cells (B). (C, D) Differential expression analysis of SO versus NO oocytes (C) and granulosa cells (D) reveals few significant genes. (E) Pseudotime analysis computed on highly variable genes in mice granulosa cells in all four experimental conditions. (F) Comparison of gene expression between NY, NO and SO oocytes using SMART-seq2. The fold change is computed between the old and young groups. Each column represents gene expression in an individual oocyte. (G) Gene expression of genes involved in cGMP pathway (left) or spindle assembly checkpoint (SAC) machinery (right) in NY, SY, NO and SO granulosa cells or oocytes, respectively.

#### Supplementary movies

**Movie S1.** Example movie for microsurgical cumulus-oocyte complex isolation, and oocyte and associated granulosa cell singularization.

#### Supplementary tables

**Table S1.** QC table for SMART-Seq2 samples (gene expression and IVF).

**Table S2.** Pairing of oocytes and granulosa cell samples used in the gene expression analyses.

**Table S3.** Cell-type-specific marker genes identified from our data.

**Table S4.** Overrepresentation analysis table for oocytes and granulosa cells in SMART-seq2.

**Table S5.** QC table for total RNAseq samples.

**Table S6.** Gene set enrichment analysis result table for human versus mouse granulosa cells comparison (only pathways that were significant in at least one species are included).

**Table S7.** Ovulation rates for naturally and superovulated young and old mice.
